# Petascale pipeline for precise alignment of images from serial section electron microscopy

**DOI:** 10.1101/2022.03.25.485816

**Authors:** Sergiy Popovych, Thomas Macrina, Nico Kemnitz, Manuel Castro, Barak Nehoran, Zhen Jia, J. Alexander Bae, Eric Mitchell, Shang Mu, Eric T. Trautman, Stephan Saalfeld, Kai Li, Sebastian Seung

## Abstract

The reconstruction of neural circuits from serial section electron microscopy (ssEM) images is being accelerated by automatic image segmentation methods. Segmentation accuracy is often limited by the preceding step of aligning 2D section images to create a 3D image stack. Precise and robust alignment in the presence of image artifacts is challenging, especially as datasets are attaining the petascale. We present a computational pipeline for aligning ssEM images with several key elements. Self-supervised convolutional nets are trained via metric learning to encode and align image pairs, and they are used to initialize iterative fine-tuning of alignment. A procedure called vector voting increases robustness to image artifacts or missing image data. For speedup the series is divided into blocks that are distributed to computational workers for alignment. The blocks are aligned to each other by composing transformations with decay, which achieves a global alignment without resorting to a time-consuming global optimization. We apply our pipeline to a whole fly brain dataset, and show improved accuracy relative to prior state of the art. We also demonstrate that our pipeline scales to a cubic millimeter of mouse visual cortex. Our pipeline is publicly available through two open source Python packages.

## Introduction

In serial section electron microscopy (ssEM), a biological sample is sliced into ultra-thin sections, which are collected and imaged at nanoscale resolution (Harris et al. 2006). When applied to brain tissue, this technique can yield a 3D image with sufficient resolution for reconstructing neural circuits by tracing neurites and identifying synapses (Briggman and Bock 2012). The technique has been scaled up to image an entire *Drosophila* brain (Zheng et al. 2018), and cubic millimeter volumes of mammalian cortex (Yin et al. 2020; MICrONS Consortium et al. 2021; Shapson-Coe et al. 2021).

Purely manual reconstruction of neural circuits from ssEM images is slow and laborious. It is faster to proofread an automated reconstruction generated via convolutional nets, as reviewed by (Lee et al. 2019). However, automated reconstruction depends on the preceding step of aligning the 2D section images into a 3D image stack. Misalignments can be the dominant cause of errors in the automated reconstructions, so it is important for the alignment to be precise and robust to image artifacts.

State-of-the-art alignment methods identify a sparse set of corresponding points between neighboring images, and use an offline solver to find image transformations that register corresponding points subject to an elastic regularizer that prevents nonsmooth transformations (Khairy, Denisov, and Saalfeld 2018; Saalfeld et al. 2012; Bock et al. 2011; Wetzel et al. 2016; Scheffer, Karsh, and Vitaladevun 2013). Correspondences may be identified by matching image patches through some kind of cross-correlation (Wetzel et al. 2016; Saalfeld et al. 2012), or through matching hand-designed features such as SIFT (Lowe 1999).

These state-of-the-art methods are inadequate in a number of ways. First, cracks and folds are common defects in serial sections (Fig. 1a). Correctly aligning pixels on both sides of a crack or fold (Fig. 1b,c) requires moving closely neighboring pixels in opposite directions. Such a nonsmooth transformation is not possible by interpolating between sparse correspondences, as done by state-of-the-art methods. And even if the density of correspondences is increased, the elastic regularizer also prevents nonsmooth transformations.

**Fig. 1.**
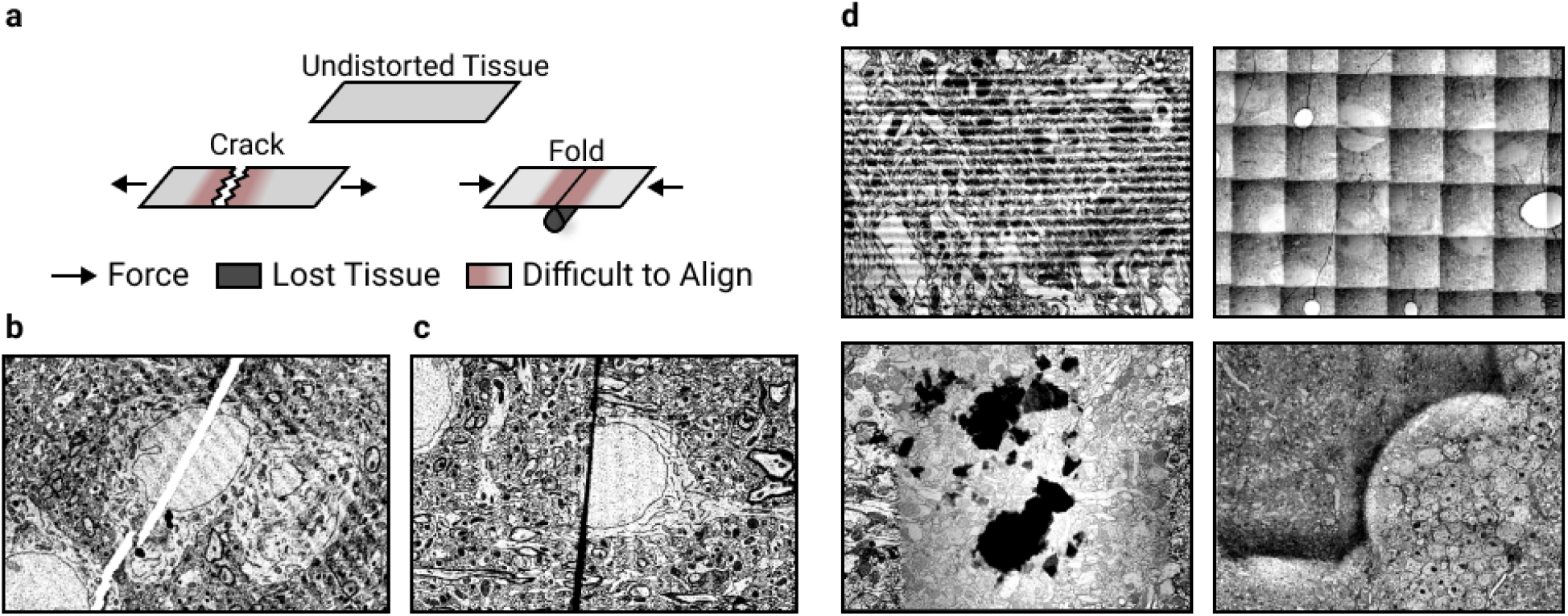
Challenges In ssEM Alignment. **a**, Mechanics of discontinuous defect creation. Discontinuous defects cause tissue loss and make neighboring areas particularly challenging to align. **b**, Example of a crack (white line). **c**, Example of a fold (black line). **d**, Examples of other artifacts: knife chatter (top left), grid pattern (top right), dust (bottom left), brightness variation (bottom right)

Second, alignment must robustly handle image artifacts. Consider a case where striped artifacts caused by knife chatter (Fig. 1d, top left) occur in two sections in a row. High similarity alignment between such a section pair can be produced in two ways: the correct alignment that aligns biological objects, and an erroneous alignment that aligns the stripe patterns. General purpose image feature extractors, such as SIFT (Lowe 1999), do not prevent such errors. While there exist methods that are able to effectively suppress EM image artifacts *after* the dataset has been aligned (Kazhdan et al. 2015), robust artifact suppression in EM images prior to alignment has not been demonstrated.

Third, even without image artifacts, state-of-the-art approaches make misalignment errors due to false correspondences. The frequency of errors is reduced by hand-designed heuristics for rejecting false correspondences (Saalfeld et al. 2012; Bock et al. 2011; Scheffer, Karsh, and Vitaladevun 2013). For a large dataset, it is difficult or impossible to set the parameters of the heuristics to remove all false correspondences, so manual intervention at the level of single sections or even single correspondences becomes necessary (Turner et al. 2020).

Fourth, the offline solver in state-of-the-art methods does not scale well to large datasets. The number of variables in the optimization increases with image volume and the complexity of the alignment transformations. We have found that the optimization can be slow because long-wavelength modes of an elastic energy tend to relax slowly. It could be possible to speed up convergence by advanced optimization techniques. Here we take a different approach, which is to completely eliminate the need for a global optimization over the entire dataset.

The above difficulties are all overcome by our new computational pipeline for alignment of ssEM images. The input to the pipeline is assumed to have been already aligned coarsely, which can be done by standard methods. The global rotations and scale changes in the input, for example, should be small. We show that our pipeline significantly improves alignment over state-of-the-art methods on the challenging whole brain female adult fly dataset (FAFB) and can align a petascale mouse cortex dataset (MICrONS Consortium et al. 2021). The pipeline demonstrates unprecedented ability to align near cracks and folds up to 10 μm wide.

Our pipeline contains several novel elements. First, we use metric learning to train convolutional nets to extract encodings of EM images, and align the images using the encodings (Fig. 5). The method of training is called Siamese Encoding and Alignment by Multiscale Learning with Self Supervision (SEAMLeSS), though there are some differences from a preliminary version of the method (Mitchell et al. 2019) that will be described later. Alignment based on the encodings is more robust, because image artifacts are suppressed by the encodings. The convolutional nets are applied in a coarse-to-fine hierarchy (Fig. 2a), which enables correction of large displacements. The use of convolutional nets to compute dense correspondences between images is now standard in computer vision (Yoo et al. 2017; Balakrishnan et al. 2019; Zhou et al. 2019; Shu et al. 2020; Nguyen-Duc et al. 2019; Jain 2017), and is compatible with nonsmooth transformations.

**Fig. 2.**
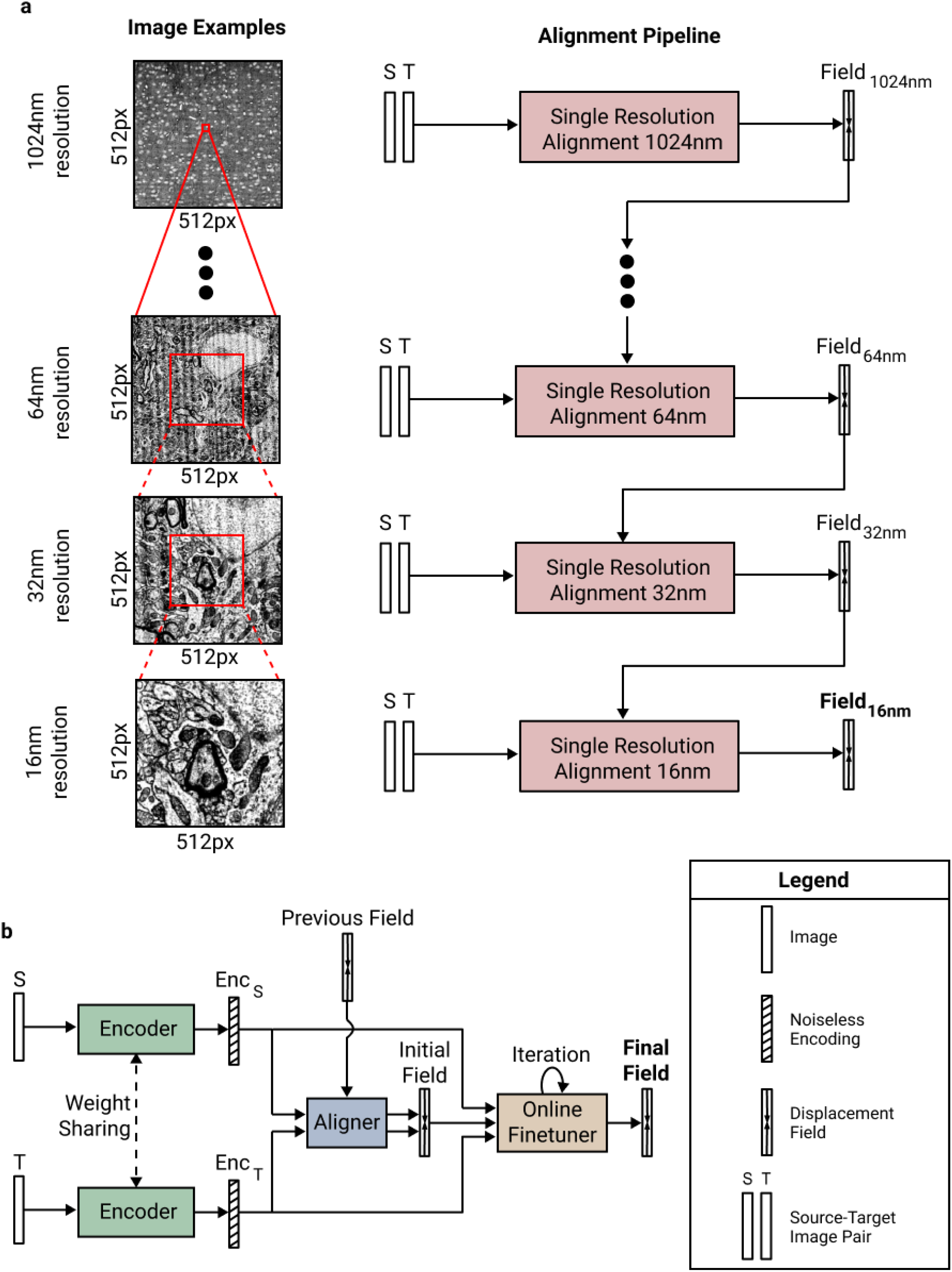
SEAMLeSS Image Pair Alignment. **a**, Coarse-to-fine multi resolution alignment. Input to each subsequent stage increases physical resolution, while keeping the same pixel resolution. **b**, Architecture of a single resolution alignment pipeline: Encoder, Aligner and Online Field Finetuner.

Second, the alignment transformation is fine-tuned by gradient descent on a cost function that is the squared difference between the image encodings plus an elastic regularizer. Gradient descent is initialized at the alignment transformation that was generated by the convolutional net. The fine-tuning achieves more precise alignment than SEAMLeSS alone, at the expense of moderately more computation. Both in SEAMLeSS training and in fine-tuning, the elastic regularizer is ignored at locations where nonsmooth transformations are required, using a convolutional net trained to detect cracks and folds.

Third, for more robust alignment, each section is aligned to several preceding sections and a procedure called vector voting is used to arrive at a consensus alignment. If the data were defect-free, it would be sufficient to align each section to the previous section. But virtually every section contains one or more regions with defects or missing data. Therefore it is helpful to also align to sections that are further back in the series. Previous elastic alignment schemes added springs between the next nearest neighbor and further sections, motivated by similar considerations (Saalfeld et al. 2012). Vector voting combines multiple alignment transformations with a smooth pixel-wise operation (Fig. 3a) that is robust to the presence of outliers.

**Fig. 3.**
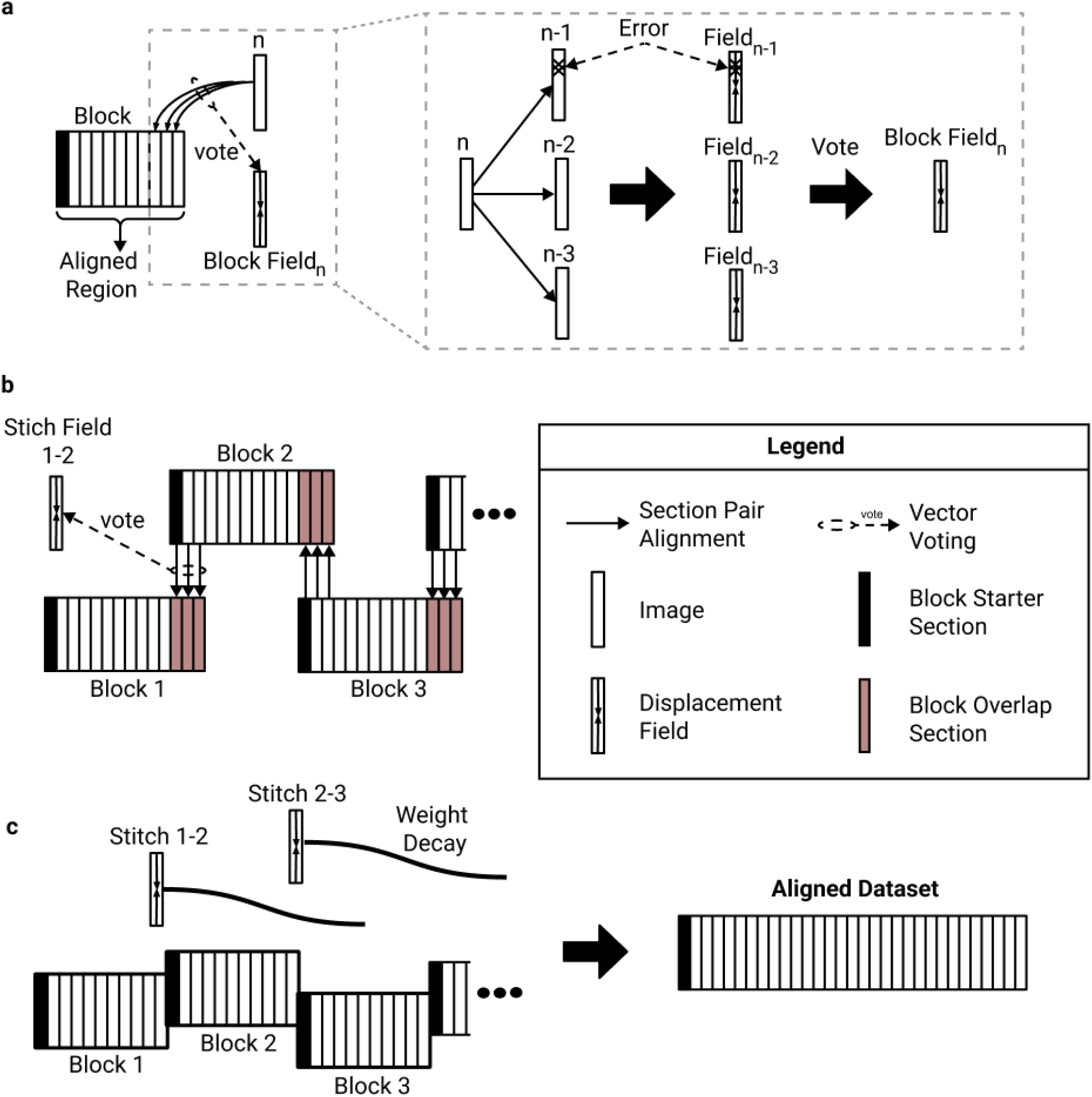
Vector Voting, Distributed Block Alignment, and Composition with Decay. **a**, Sequential alignment with vector voting. During block alignment sections are aligned sequentially to the already aligned portion of the stack. Vector voting is performed by aligning the source image to multiple target images, and passing the resulting fields through a pixel-wise smooth median computation. **b**, Producing block stitching fields. To produce global alignment, the dataset is broken into overlapping blocks, which are aligned independently. The overlapping regions are aligned to each other to produce a stitching field between each two neighboring blocks. **c**, Block stitching. The blocks are stitched together by applying each stitching to successive sections, while reducing its influence with a distance based weight decay.

Fourth, we divide the entire series of sections into blocks of contiguous sections, and sequentially align the sections in each block. The blocks are distributed over multiple computational workers, speeding up the computation.

Fifth, we have to combine the aligned blocks into a single global alignment. Naively, this could be done by aligning the start of each block with the end of the previous block, and then composing these transformations. If there are many blocks, however, the composition of transformations could result in highly distorted sections later in the series. This was previously solved by a global relaxation of a spring mesh that extends across the entire series of sections (Saalfeld et al. 2012). Here we avoid any global relaxation, which would be computationally expensive for petascale datasets with dense high-resolution transformations. Instead, we compose transformations, but force each transformation to decay in time/depth and in spatial frequency. We call this approach Alignment of Blocks and Composition with Decay (ABCD).

We release two open-source packages, *metroEM* and *corgie*, that together implement our computational pipeline. *metroEM* implements SEAMLeSS training and inference for image pair alignment. All the multi-resolution Encoder/Aligner pairs (Fig. 2a) can be trained with a single *metroEM* command. *corgie* implements large scale global alignment, including block alignment, stitching, and vector voting. *corgie* is able to handle images that do not fit into single machine memory, provides intuitive tools for distributing tasks to cloud and cluster workers, and can use both *metroEM*-based or any other user supplied method for image pair alignment. Both packages are implemented using the python (3.6+) programming language and can be installed with the *pip* package-management system. The released tools enable training of the convolutional nets in our pipeline and application of the convolutional nets to align petascale datasets using cloud Kubernetes or SLURM clusters.

## Results

### SEAMLeSS Image Pair Alignment

Given a source image and a target image, the task of image pair alignment is to produce a transformation that aligns the source to the target. Transformations are represented as displacement fields, which assign a 2D displacement vector to each pixel (formal definition in Methods). SEAMLeSS works at multiple resolution scales in a coarse-to-fine manner (Bajcsy and Kovačič 1989; Ranjan and Black 2016; Wetzel et al. 2016) (Fig. 2a). At each resolution, there is an Encoder net, Aligner net, and Online Field Finetuner (Fig. 2b). The Encoder is applied to both source and target images (Bromley et al. 1993), and the resulting encodings are the inputs to the Aligner. The Aligner computes a residual displacement field, which is composed with the estimate of the displacement field from a coarser resolution, to align the source to the target. The Field Finetuner refines the initial field produced by Aligner through gradient descent to minimize the difference between the image encodings after alignment while keeping the transformation smooth in the areas without defects (Broit 1981). A separate instance of Encoder/Aligner pair is trained for each resolution level.

### SEAMLeSS Training

The training dataset for Aligner and Encoder nets consists of neighboring image pairs from an unaligned EM stack. Both Aligner and Encoder are trained in a self-supervised manner, meaning that no alignment ground truth is required for training. Training is performed sequentially for resolutions in a coarse-to-fine manner.

Training proceeds in two stages. During the first stage, the training loss is based on the squared difference between the images after alignment by the Aligner net, plus an elastic regularizer. (See Methods for precise description of the loss function.) Since nonsmooth transformations are required at cracks and folds, these defects are detected by another convolutional net, and the regularizer is zeroed out at these locations. The first training stage continues until the loss plateaus, and results in an Encoder/Aligner pair that can align tissue in most cases but is susceptible to errors due to image defects.

In the second stage of training, the training loss is based on the squared difference between the encodings (not the images) after alignment by the Aligner net. This can be viewed as metric learning, if the Aligner net is regarded as part of the (trainable) distance function. In other words, the distance function is the Euclidean distance between the encodings (plus an elastic regularizer) after alignment by the Aligner net. Then the goal of the Encoder is to produce encodings that are similar for images that can be aligned, and dissimilar for images that cannot be aligned. The goal of the optimization is to produce image encodings and alignment fields such that the pixel-wise difference between encodings is high before alignment, but low after alignment. Under such training, the Encoder will learn to suppress image artifacts while preserving information about the biological objects. Image artifacts must be suppressed to achieve low pixel-wise difference between aligned encodings because image variation due to artifacts is not correlated across the same pixel positions in neighboring aligned sections. At the same time, highlighting biological objects will increase the difference between unaligned encodings. In order to satisfy both training goals, the Encoder ends up suppressing image artifacts and highlighting biological objects.

It is desirable for the encodings to include information about each pixel in the image, as opposed to sparsely representing selected features. With dense encodings, it is possible to precisely correct discontinuous defects based on encoding similarity. We introduce additional training constraints to encourage encodings to be dense as described in Methods. At the end of the second training stage, the Encoder is able to suppress artifacts in EM images.

After a given resolution scale Aligner/Encoder pair has finished training, it is used to process the whole training dataset to produce displacement fields that will be used for subsequent resolution scales training. Training of multi-resolution hierarchies of Aligner/Encoder pairs is implemented in the metroEM package released with this work.

### Online Field Finetuner

Online Field Finetuner refines the initial displacement field through gradient descent. The goal of the optimization process is to maximize similarity between image encodings after alignment, while penalizing solutions that produce unnatural distortions.

Finetuning by gradient descent is facilitated by using the encodings. Using the raw images instead may cause catastrophic errors because image artifacts are often high contrast, and may end up being the features that are aligned (Fig. 1d). For example, alignment can be corrupted by the parallel stripes of knife chatter (Fig. 1d, top left).

### Vector Voting

In principle, one could align an entire series of sections by applying SEAMLeSS to align each new section to the previously aligned section. This sequential alignment procedure is well-known (Tasdizen et al. 2010) and effective if each alignment step is perfect, but lacks robustness to occasional misalignments. These may happen near image artifacts, and are unavoidable in regions where the image data is entirely missing. To increase robustness, we instead align the source image to several last sections of the already aligned portion (Fig. 3a) and apply a pixel-wise smooth median computation over the resulting displacement fields. This Vector Voting operation discards the outlier values for each pixel location of the field, ensuring that only errors that occur at the same pixel location in several neighboring sections will propagate through the block (Methods). Aligning source sections to *n* target sections is referred to as *n*-way Vector Voting, where *n* is referred to as voting distance. For *n*-way Vector Voting with odd *n*, only the errors that occur in *(n + 1) / 2* out of *n* consecutive sections will be propagated through the block. Increasing voting distance improves error resilience at the cost of increasing the computational cost of alignment.

In alignment methods based on relaxation of a spring mesh (Saalfeld et al. 2012), springs are sometimes added between next nearest and further neighbor sections, in addition to the springs between nearest neighbor sections. This is a kind of voting, and tends to reduce the magnitude of a misalignment due to a false correspondence but also spreads the misalignment over a larger area. The robustness delivered by Vector Voting should be superior because the effects of outliers are suppressed almost completely.

### Distributed Alignment of Blocks

In order to speed up the computation, we partition the entire series along the cutting dimension into overlapping blocks of contiguous sections. Each block is sequentially aligned with Vector Voting, and the blocks are distributed over independent computational workers.

Vector Voting is not possible for the first *(n - 1) / 2* sections in a series, which increases the likelihood of error across that region. Therefore adjacent blocks are chosen to overlap by *(n - 1) / 2* sections, and the first *(n - 1) / 2* sections of each block are discarded before combining the blocks to create a global alignment (Fig. 3b), as will be discussed next.

### Composition with Decay

After all the independent blocks are aligned, they should be stitched together into a single globally aligned stack. We define the stitch field between block *N* and block *N+1* as that obtained by aligning the overlapping section region of block *N+1* to the same sections in block *N* and performing vector voting operations across the obtained fields (Fig. 3b). The stitch fields can be used to create a global alignment as follows. For each block, the stitch fields from the preceding blocks are composed to create an accumulated transformation that is applied to all sections in the block (Fig. 3c). This straightforward procedure can result in good alignment, in the sense that neurites follow smooth paths through the stack. However, the composition of many stitch fields generally results in a highly distorted transformation (Fig. 4g). This is because the composition is not limited by the elastic regularizer, even though the individual stitch fields are governed by the elastic regularizer.

**Fig 4.**
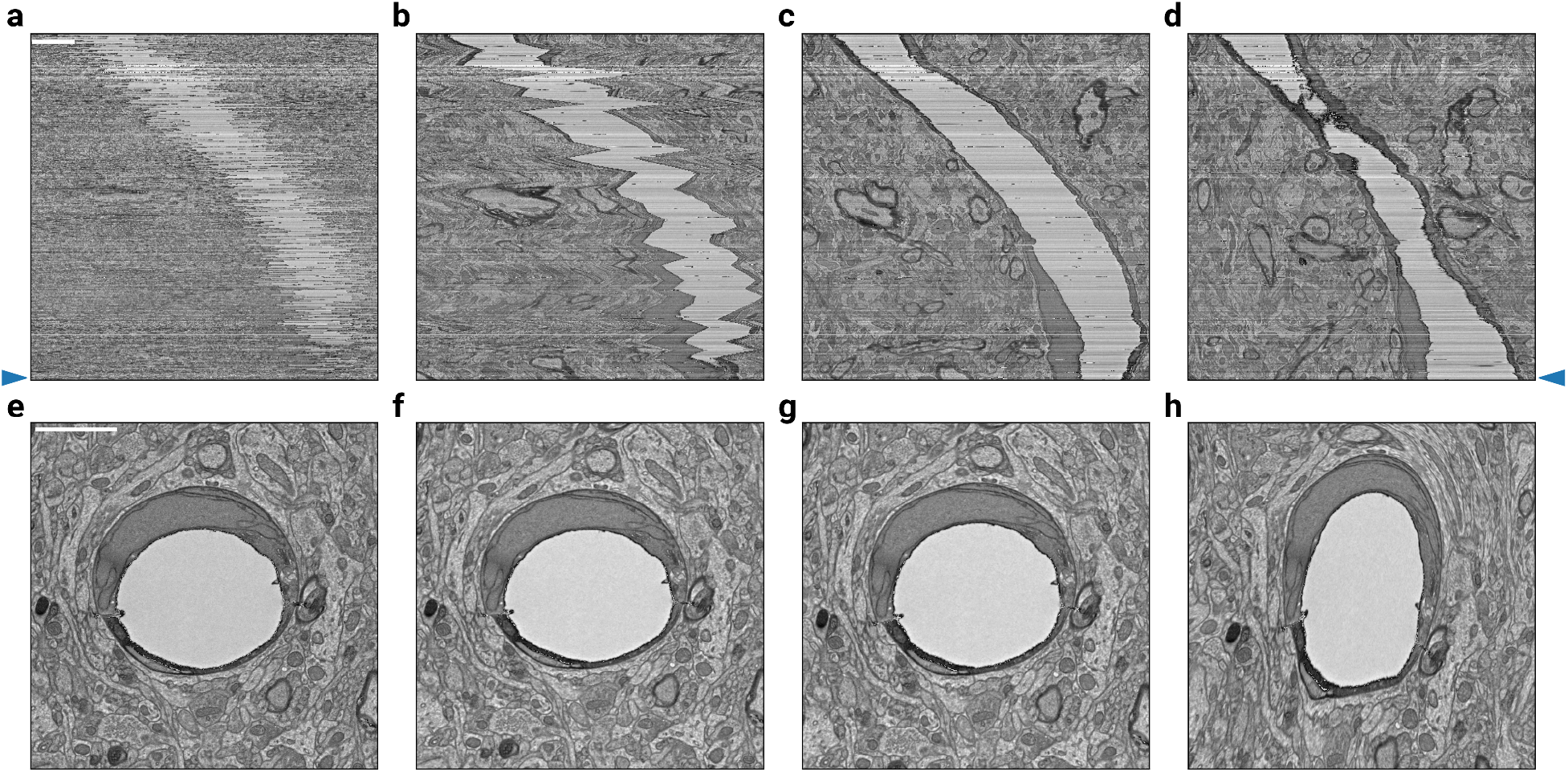
Alignment of Blocks and Composition with Decay. A stack of 450 mouse cortex sections (MICrONS Consortium et al. 2021) was block aligned (10 sections per block), then globally aligned by composing the linearly decayed stitching fields between blocks with varying decay distances. **a-d**, The rough aligned sections, the aligned sections with decay distances of 10, 100, and 500 sections. The vertical axis is orthogonal to the cutting plane, while the horizontal axis is in the plane. **e-h**, The 447th section of the sample above (see blue arrow along sides of a and d), approximately centered on the same blood vessel to demonstrate the distortion that results from composing many fields together without decay. The horizontal axes of the top and bottom row are the same. Scale bars are 2 μm.

Therefore we modify the procedure as follows. When applying the composition of stitch fields to a section, each stitch field is forced to decay by an amount that depends on how far it was back in depth (Methods). The number of sections that each stitch field influences is referred to as *decay distance*. Using very large decay distances may result in large distortions due to composition of many transformations (Fig. 4h), whereas very small decay distances may result in drifts between sections (Fig. 4b, f).

Alignment of Blocks and Composition with Decay (ABCD) avoids a global optimization with an offline solver, which becomes computationally expensive with dense high-resolution fields (Khairy, Denisov, and Saalfeld 2018; Saalfeld et al. 2012),

### Whole fly brain alignment

We evaluate our alignment pipeline by applying it to the full adult fly brain (FAFB) dataset (Zheng et al. 2018). As a baseline, we first consider the original alignment (v14), which was created using a piecewise affine approach (Khairy, Denisov, and Saalfeld 2018). To evaluate alignment accuracy, we adopted the Chunked Pearson Correlation framework (Möller, Garcia, and Posch 2007). The evaluation was performed on 1000 full sections of v14 at 32 nm pixel resolution. Each section was divided into 64×64 chunks, roughly 2 µm across at this resolution. Chunks containing non-tissue (blank) areas were excluded. Chunks containing cracks or folds were also excluded; chunks near such discontinuous defects were included. We computed the Pearson correlation for chunk pairs in neighboring sections.

We found that 4.23% of v14 chunks had CPC values less than a threshold value of 0.25 (Fig. 5a, b). The CPC tends to be low at misalignments (Fig. 5c). However, the CPC can be low for reasons other than misalignment, such as image artifacts or genuine changes in image content across neighboring sections (Fig. 4d). Therefore 300 chunks with CPC less than 0.25 were selected at random, and examined by a human annotator. Based on the results, we estimate that 3.31% of v14 contains a genuine misalignment with CPC less than 0.25 (Fig. 5b).

**Fig. 5.**
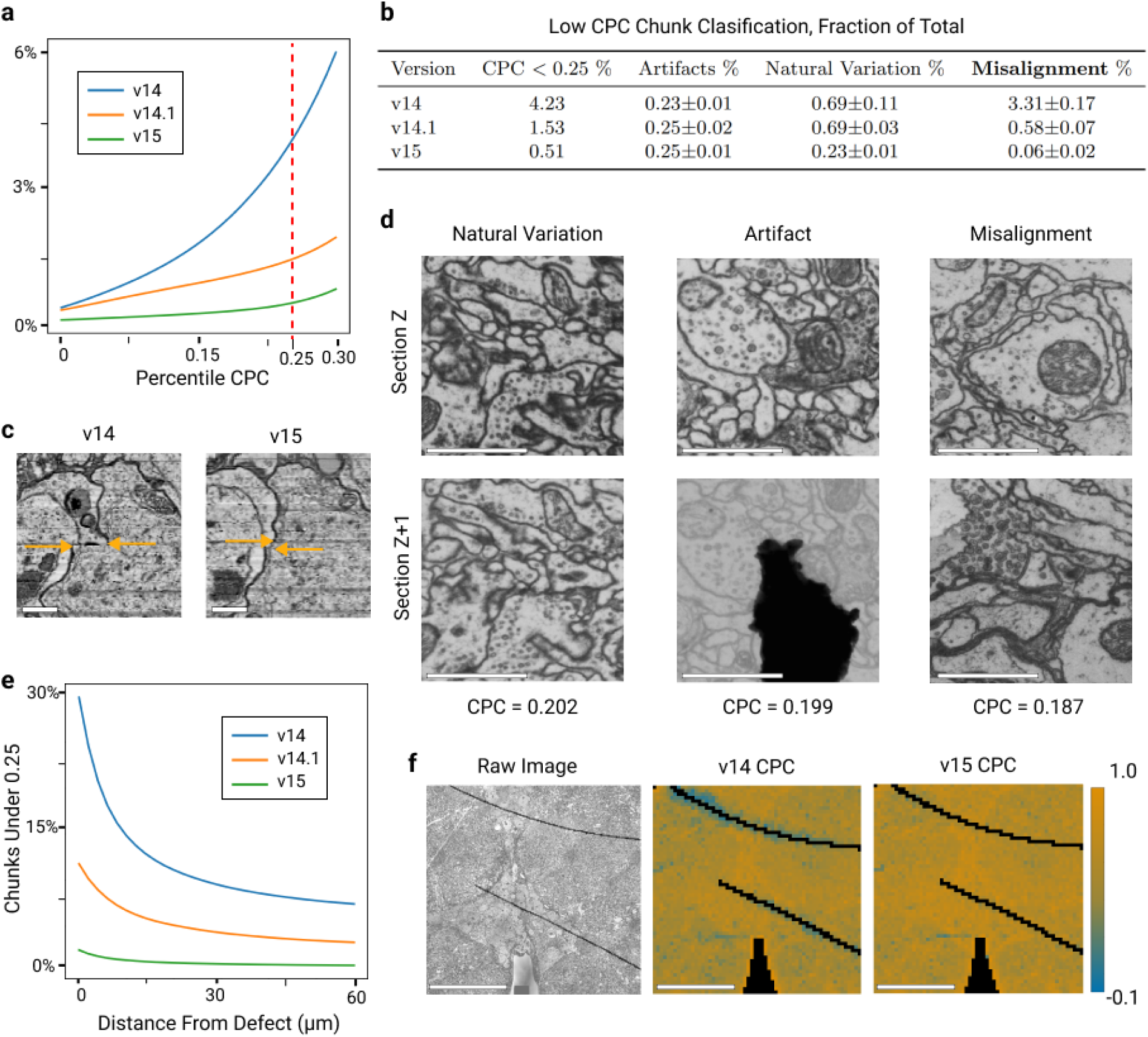
FAFB Alignment Quality. A new v15 alignment of the FAFB dataset improves over v14.1 (Dorkenwald et al. 2021), which in turn improves over the original v14 (Zheng et al. 2018). Accuracy is quantified using the Chunked Pearson Correlation (CPC), which tends to be high at well-aligned locations. The evaluation was conducted for a range of 1000 sections. **a**, CPC percentile decreases with progressive improvement of the three alignment versions. Chunks with low CPC (<0.25, dashed red line) are just 0.51% of v15. CPC was computed for (2048 nm)^2^ chunks at 32 nm pixel resolution, and non-tissue chunks and chunks that include a discontinuous artifact were excluded. **b**, Estimated misalignment rate based on manual annotation of low CPC chunks. v15 improves over v14 by two orders of magnitude (rightmost column). CPC may be low for reasons other than misalignment, so 100 chunks with CPC < 0.25 were randomly sampled from each alignment version, misalignments were identified by a human expert, and the results were extrapolated to the entire image stack. **c**, Example misalignment in v14 corrected in v15. Arrows indicate cell body boundary position, and the horizontal space between the arrow heads indicates misalignment. Horizontal and vertical axes are parallel and perpendicular to the sections, respectively. Scale bar 1 μm. **d**, Examples of low CPC (<0.25) chunks caused by natural variation of image content, image artifact and misalignment, the categories that were manually annotated in (b). Note that a misalignment of a large magnitude was chosen for visual clarity. Scale bar 1 μm. **e**, Percentage of low CPC (<0.25) chunks as a function of distance from a discontinuous defect. In v15, such defects cause little increase in low CPC chunks. **f**, Single section CPC heatmap comparison between v14 and v15. Black corresponds to non-tissue and discontinuous defect chunks, which are ignored in (a). Scale bar 50 μm.

While the great majority of v14 is well-aligned, the misaligned portion poses challenges for automated segmentation. For example, inspection of an automated segmentation of v14 (Li et al. 2019) that is publicly available reveals that v14 misalignments generally lead to errors in reconstructed neurons.

Therefore we were motivated to create a new v15 alignment of FAFB using the pipeline presented in this paper. Only 0.51% of v15 has CPC less than 0.25 (Fig. 5a, b). Based on random sampling of chunks and human annotation, we estimate that just 0.06% of v15 contains a genuine misalignment with CPC less than 0.25. This is almost two orders of magnitude better than v14.

Not all of the v15 improvement can be attributed to alignment, because the 2D section images for v15 were reworked to remove montaging errors in v14. To isolate the improvements due to alignment rather than montaging, we evaluated another FAFB alignment (v14.1) that was previously released (Dorkenwald et al. 2021). v14.1 was created by applying a preliminary version of our pipeline (Methods) to v14 images, so that montaging errors in v14 were “baked in” and could not be corrected. We estimate that 0.58% of v14.1 contains a genuine misalignment with CPC less than 0.25. This is almost 6× better than v14, suggesting that some of the improvement in v15 is due to alignment alone. This is an underestimate of the impact, because alignment and montaging interact in v14; many of the v14 montaging errors are caused by false correspondences arising in alignment.

### Petascale alignment of mouse cortex

A preliminary version of our pipeline, similar to the one used for the v14.1 FAFB alignment, was applied to two mouse cortex volumes amounting to over 1.1 mm^3^. The largest mouse cortex dataset (MICrONS Consortium et al. 2021) was composed of 19,939 square millimeter sections imaged at 4 nm resolution (Fig. S2). Numerous cracks and folds occurred in every section, posing challenges more severe than in FAFB (Fig. S3). The discontinuous distortion introduced by cracks and folds ranged up to roughly 10 μm, with the cross-sections of neurites nearly 1000× smaller than the distortion. Even our preliminary pipeline was able to precisely correct such large discontinuous distortions, ensuring that neurites at the boundary of the defect were still well-aligned. This led to a successful automated reconstruction (Macrina et al. 2021). The alignment took 230 hours on a cluster of preemptible NVIDIA T4 GPUs on Google Cloud. The cluster size fluctuated due to hardware availability constraints, with the number of active GPUs averaging at 1200.

## Discussion

Our pipeline computes dense displacement fields, rather than the sparse correspondences of traditional approaches (Saalfeld et al. 2012; Khairy, Denisov, and Saalfeld 2018). This enables the representation of nonsmooth transformations, which are necessary for accurate alignment near cracks and folds. Displacement fields are initially predicted by the SEAMLeSS convolutional nets, and then improved by the Online Field Finetuner. This combination is more reliable at avoiding false correspondences than traditional approaches based on Block Matching or SIFT features, as shown by our evaluation on the FAFB dataset.

The rate of false correspondences can be reduced in traditional approaches by various postprocessing schemes (Saalfeld et al. 2012), but even a low rate amounts to a large absolute number of false correspondences in a sufficiently large dataset, as in the FAFB v14 alignment. For a previous terascale alignment, we eliminated the remaining false correspondences by manual intervention (Turner et al. 2020), but the required labor would be prohibitive at the petascale. Our approach is to drastically reduce the rate of false correspondences by training the SEAMLeSS nets.

SEAMLeSS nets were used for the FAFB v14.1 alignment (Dorkenwald et al. 2021) in a one-shot manner, without subsequent iterative improvement by the Online Field Finetuner (Mitchell et al. 2019). When creating v15, we found that the Finetuner is helpful for improving the accuracy of alignment, especially near folds and cracks. Also, the Finetuner reduces the accuracy requirements for the SEAMLeSS nets, making training them easier.

Conversely, one can imagine eliminating SEAMLeSS from the pipeline, and using the Online Field Finetuner only, but this would lower accuracy for two reasons. By providing a good initial guess for the displacement field, the SEAMLeSS nets prevent the Finetuner from getting trapped in a bad local optimum (Mitchell et al. 2019). Furthermore, the SEAMLeSS Encoder suppresses image artifacts (Fig. 6), which otherwise can cause misalignments to become global optima.

**Fig. 6.**
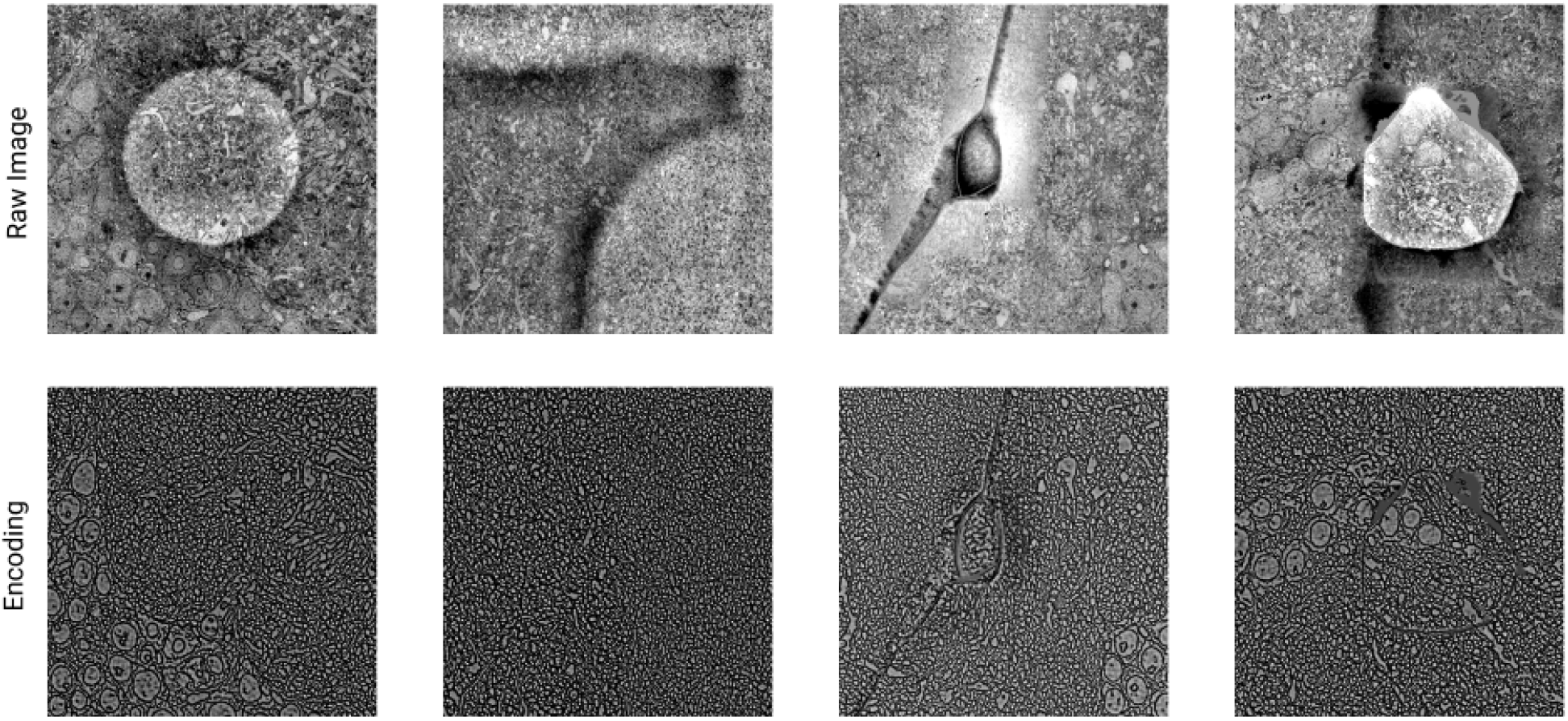
Encodings suppress artifacts. SEAMLeSS is able to produce dense Encodings that suppress many artifacts present in ssEM images. As a result, Encodings are more suitable than images as inputs to the Online Field Finetuner.

Our ABCD procedure provides a computationally efficient alternative to traditional approaches involving global optimizations over all sections in an image stack (Saalfeld et al. 2012). All ABCD computations are local, involving neighboring sections only. Within each block of sections, a section is aligned to the previous *n* sections considered by the Vector Voting procedure. When stitching blocks together, blocks do not interact with other blocks beyond the decay distance. Such locality enables graceful extension to petascale datasets, and the blocks can be processed in parallel by hundreds of distributed workers.

In the preceding, we have emphasized the replacement of traditional computer vision by our approach. In reality, the approaches are complementary and can be combined. This is already assumed by the present paper, as our pipeline is applied only after the images have been coarsely aligned using traditional computer vision approaches (Methods). Although our approach does not require a global spring mesh extending over all sections, an elastic regularizer is used for 2D patches during SEAMLeSS training and by the Online Field Finetuner at runtime. One can easily imagine other hybrid methods. For example, Block Matching should be more robust to image artifacts if applied to our SEAMLeSS encodings rather than to raw images or linearly filtered images. Our ABCD procedure, including Vector Voting, could be applied to transformations computed by Block Matching rather than SEAMLeSS. These examples of hybrid methods are hypothetical, and are left for future work.

## Acknowledgements

Research was supported by the Intelligence Advanced Research Projects Activity (IARPA) via Department of Interior/Interior Business Center (DoI/IBC) contract numbers D16PC00003, D16PC00004, and D16PC0005. The U.S. Government is authorized to reproduce and distribute reprints for Governmental purposes notwithstanding any copyright annotation thereon. HSS also acknowledges support from NIH/NINDS U19 NS104648, NIH/NEI R01 EY027036, NIH/NIMH U01 MH114824, NIH/NINDS R01NS104926, NIH/NIMH RF1MH117815, NIH/NIMH RF1MH123400 and the Mathers Foundation, as well as assistance from Google, Amazon, and Intel. These companies had no influence on the research. We are grateful for support with FAFB imagery from D. Bock through NIH/NINDS RF1MH120679, and software administrative support provided by Tom Kazimiers (Kazmos GmbH) and Eric Perlman (Yikes LLC). We are grateful for support with mouse cortex imagery from N. da Costa, R. Torres, G. Mahalingam, R.C. Reid. We thank A. Burke, J. Gager, J. Hebditch, S. Koolman, M. Moore, S. Morejohn, B. Silverman, K. Willie, R. Willie for their image analyses, S-C. Yu for managing image analysis, G. McGrath for computer system administration, and M. Husseini and L. and J. Jackel for project administration. We are grateful to J. Maitin-Shepard for making Neuroglancer freely available. Disclaimer: The views and conclusions contained herein are those of the authors and should not be interpreted as necessarily representing the official policies or endorsements, either expressed or implied, of IARPA, DoI/IBC, or the U.S. Government.

## Author contributions

### FAFB v15

ET and SS revised and rigidly aligned montages from FAFB v14. AB trained crack and fold detection models. NK trained tissue detection model. NK coarse aligned using aligner trained by SP. NK fine aligned using aligners trained by SP, TM.

### Mouse cortex demonstration

Demonstration was aligned by SP using aligners trained by SP.

### metroem development

SP developed. TM added methods to improve usability.

### corgie development

SP developed. TM added methods for vector voting and ABCD. NK, MC added features and fixed bugs.

### Preliminary pipeline development

Preliminary version of SEAMLeSS developed by EM. Preliminary pipeline codebase architected by SP, with contributions from TM, NK, MC, BN, ZJ, AB, SM. KL consulted on development.

### Mouse cortex in MICrONS release

SP, MC, NK, TM used the preliminary pipeline to align MICrONS mouse cortex samples.

### Manuscript

SP, TM, HSS wrote the manuscript.

### Leadership

HSS led the project.

## Competing Interest

TM and HSS disclose financial interests in Zetta AI LLC.

## Methods

### Female Adult Fly Brain (FAFB)

#### v14.1

The v14.1 alignment was produced using a preliminary version of the pipeline described in this paper applied to the v14 alignment of the FAFB dataset (Zheng et al., 2018). At this writing, the v14.1 alignment is being used in FlyWire, an online community for proofreading an automated segmentation of FAFB (Dorkenwald et al., 2022).

The preliminary pipeline did not include an Encoder or Online Field Finetuner. The pipeline relied only on a preliminary version of SEAMLeSS, as described in a previous preprint (Mitchell et al., 2019). The loss function for self-supervised learning was based on the difference between source and target images rather than encodings after alignment, therefore the training procedure is not considered deep metric learning. During training of the Aligner, precomputed defect masks were used to relax the smoothness penalty, but no defect masks were provided at inference time. In the current pipeline, precomputed defect masks are included as an input to the Aligner. The preliminary pipeline used Vector Voting with a voting distance that was manually set for each section. The voting distance was typically 5, with some small regions set to 7 or 9, allowing additional computation to be spent for specific sections that needed to contend with long stretches of missing data.

#### v15

Many v14.1 problems were the result of errors in v14 that could not be undone by our alignment pipeline. Every section image is composed of multiple image tiles, which are combined by a process known as montaging, stitching, or mosaicking. Montages in v14 were generated using SIFT feature matches, geometric consensus filtering and global optimization of per-tile affine transformations (Saalfeld et al., 2012; Khairy et al., 2018; Zheng et al., 2018). Many montages in v14 have substantial stitching errors. To eliminate these errors, we attempted to produce a clean set of montages that would serve as the starting point for our alignment pipeline.

We enhanced the EM_Aligner montage diagnostic tool^1^ and found that 2,871 out of 7,050 montages had substantial point-match residuals (≥ 1% of all tile pairs having an error ≥ 10 px, the cellular membrane thickness at 4 nm resolution is ≈ 5 px). Of the remaining 4,179 montages 4,074 consisted of disconnected ‘islands’ that were improperly interleaving which corrupted the montages.

We improved the quality of point matches for stitching by re-parameterizing both feature extraction and geometric consensus filters. We used five separate sets of parameters^2^ and iterated through them sequentially until a sufficient number of matches were derived. This iterative approach allowed us to generate most matches quickly and spend more compute time only where necessary. Remaining unmatched tile pairs were manually reviewed and we tried different custom match derivation parameter sets to connect them. Often, the most difficult to match pairs contained large areas of featureless resin in their overlapping region. We further improved the global optimization of per-tile transformations. We used the EM_aligner solver (Khairy et al., 2018) that was used for the v14 reconstruction and later the Python implementation BigFeta^3^. For montages where these solvers did not produce satisfying results, we used the iterative TrakEM2 solver (Saalfeld et al., 2012)^4^ that enables rigid regularization of per-tile affine transformations but is significantly slower. This was primarily necessary for the 295 sections that contained images from multiple acquisitions.

We separated the disconnected islands in all 7,050 sections resulting in 48,121 sub-sections. These sub-sections were then rigidly registered to the corresponding sub-section in the FAFB v14 aligned data set (Zheng et al., 2018). Islands that were not present in the v14 data set, were placed based on stage coordinates of the microscope. This provided a good starting location for the cross section alignment process, essentially using v14 results to roughly align the entire volume.

As in v14, we used the Distributed Multi-Grid library (Kazhdan and Burns, 2015) to intensity correct all data and corrected a number of previously undetected mistakes. With the updated montaging pipeline, the FAFB montage series consisting of 14,442,143 tiles can be generated in ≈ 200,000 CPU hours.

The rigid registration of each island to the v14 data set still left scale changes and other non-rigid deformations in the stack. Some of the non-rigid deformations were obviously non-affine. For example, it was common for one portion of the section to rotate while another portion counter-rotated. We decided not to use the microCT image of the dataset before serial sectioning as a target to improve on the non-affine deformations at this stage, because the microCT image did not contain the entire dataset in its field of view.

We observed that sections in ranges 1–1500 and 5850–7060 frequently contained scale jumps, but no section-wide, non-affine deformations. Conversely, range 1500–5850 frequently contained section-wide counter-rotating deformations, but no sudden change in scale. To improve upon the rigid registration for both cases, each island was aligned to its corresponding v14 section using two different approaches: (1) An affine transformation estimate followed by an Online Field Finetuner with a strong rigidness constraint, operating on 512 nm encodings, as well as (2) two chained Online Field Finetuners with a much lower rigidness constraint, operating at 8192 nm and 512 nm, respectively.

While the chained Finetuner approach was powerful enough to correct even extreme rotational deformations, it inevitably also attempted to mimic errors present in the v14 image stack, such as large scale pinches and shifts caused by montaging errors. Therefore, we opted to always choose the near-affine approach for the two identified outer ranges that were manually confirmed to not contain severe non-affine global deformations. For the inner range, the decision was made for each section individually by subdividing the section into smaller blocks, calculating the Multiscale-SSIM (Wang et al., 2003) between both v15 alignment approaches and the v14 target for each block, and picking the approach for which the first decile of blocks had the higher similarity.

Next, the combined island images were aligned using 4 Aligner/Encoder pairs at 512 nm, 128 nm, 64 nm, 32 nm resolutions. Each Aligner/Encoder pair was trained for 600 epochs (100 epochs first stage + 500 epochs second stage). Alignment was performed in blocks of 25 sections with 3-way vector voting and 100 section decay distance. Alignment for the whole FAFB (7052 sections) performed on a cluster with 128 Ampere class NVIDIA GPUs and was completed in 240 hours.

#### Identification of misalignments

Classification of locations with CPC under 0.25 was performed by expert human annotators. The cause of low correlation at each location was assigned one of three classes – Visual Artifact, Natural Variation, and Misalignment. For each alignment version, we randomly sampled 300 locations out of the set of all chunks with CPC under 0.25. Classification was performed according to the following criteria:

1. Location where one of the two neighboring images included artifacts or tissue corruption that prevents clear tracing of biological object are classified as Visual Artifact.
2. Locations with relative motion sufficient to break biological objects which is consistent with motion of the same objects in preceding and following sections, and where the motion corresponds to small areas or naturally fast moving structures, such as axon bundles and tissue boundary, are classified as Natural Variation.
3. Locations with relative motion sufficient to break biological objects which is not consistent with motion of the same objects in the preceding and following sections are classified as Misalignment.
4. Locations with relative motion sufficient to break of biological objects which is consistent which with motion of the same objects in preceding and following sections, and where the motion corresponds to a large area of not naturally fast moving structures are classified as Misalignments.
5. Locations that do not qualify for criteria 1-4 and contain visual artifacts such as contrast and brightness variation are classified as Visual Artifact.
6. Locations that do not qualify for criteria 1-5 are classified as Natural Variation.

Proofreading annotations for each location along with the CPC maps and random seeds used during sampling are included with this submission.

### Petascale mouse cortex

Serial section EM images of a cubic millimeter volume of mouse cortex were acquired at the Allen Institute (Yin et al., 2020; MICrONS Consortium et al., 2021). The images were aligned using a preliminary version of our pipeline.

#### Coarse Alignment

The connectomics team at the Allen Institute for Brain Sciences stitched and aligned the image stack to displacements within 20 µm using their ASAP framework (Mahalingam et al., 2021).

The coarse alignment was further refined using an Encoder trained to optimize patch matching correlograms for 1024 nm data as described in (Buniatyan et al., 2020). The Aligner was a blockmatching method that correlated 256 px patches of the source image to 358 px patches of the target image on a Cartesian grid of 96 px. The dataset was divided into blocks of 100 sections, sequentially aligned with 5-way Vector Voting, and stitched using a decay distance of 300.

#### Fine Alignment

This preliminary version of the pipeline did not include an Encoder, but it did make use of the Online Field Finetuner. The blockmatched dataset was further aligned using an Aligner trained on 1024 nm data, and that output was again aligned by an Aligner trained on 64 nm data. This approach of aligning the entire dataset at a lower resolution before proceeding to higher resolutions has been replaced in the current pipeline with multi-resolution Image Pair Alignment.

The input to the 1024 nm Aligner was the output of the Encoder used during coarse alignment. The input to the 64 nm Aligner was raw images, with no Encoder. The 64 nm Aligner was trained on a curriculum of example images containing manually curated image artifacts.

### Current Pipeline Methods

The current pipeline was used to produce the v15 alignment of the FAFB dataset, as well as the alignment of a small sample of the petascale mouse cortex dataset that was previously aligned using a preliminary version of this pipeline (MICrONS Consortium et al., 2021).

#### Semantic Segmentation

Samples of manually annotated cracks and folds in cutouts of 1024 × 1024 px at 64 nm resolution, were collected from the original data and were used to supervise training of separate UNets as described in (Macrina et al., 2021). A tissue segmentation model was trained in similar fashion. Segmentations were downsampled with min pooling, to overestimate non-tissue regions at lower resolutions.

#### Image Preprocessing

Prior to alignment training and inference, the ssEM image stack is downsampled, masked, and normalized. First, the original images are downsampled through average pooling to produce a hierarchy of resolutions ranging up to 4096 nm. Next, discontinuous image defect masks and resin masks are produced by specialized convolutional networks. Salient visual features of cracks, folds, and resin make detection a relatively simple task. Masks are typically detected at 96 nm resolution. After the masks are produced, they’re downsampled to lower resolutions through max pooling. At each resolution, ssEM image pixels marked by the masks are zeroed-out, thus removing them from the image. Finally, the images are normalized at each resolution. Zero valued pixels, meaning pixels corresponding to empty space or the masked regions, are ignored during normalization. This means that mean and variance values used during normalization are computed based on values of non-zero pixels only, and zero valued pixels remain zero valued after the normalization. The final normalized stack of multi-resolution images with zeroed-out defect and resin regions is used during all of the alignment steps.

#### Transforming Images With Displacement Fields

Due to the large anisotropy in serial sectioning data, we consider the data as a series of 2D images,*M* : ℛ^2^ → ℛ. We define a displacement field, ***F*** : ℛ^2^ → ℛ^2^, in the transformed space, 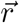, with a displacement, 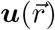, that indicates a position in the initial space,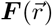, such that

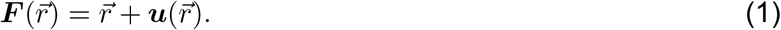

An image *M* is transformed by a displacement field by sampling,

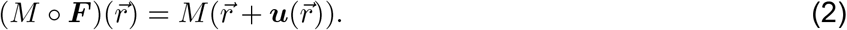

The aim of alignment is to find a displacement field that will transform each section to remove distortion. Displacement fields, 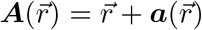 and 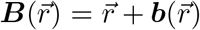 can similarly be transformed or “composed”, such that

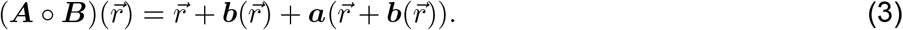

We used bilinear interpolation kernels to implement composition and sampling discretely (Jaderberg et al., 2015). We may leave off the indexing with 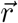 for readability and write these transformations as *M ◦* ***F*** or ***A*** *◦* ***B***.

### Loss Masking

As described in Image Preprocessing section, locations in the image that are non-tissue are set to zero. It is desirable to ignore both the similarity and the deformation loss components at such locations in the image. Given an image *M*, we define a non-zero coordinate set *T* as

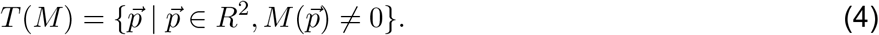

We will refer to *T (M)* as a set of tissue coordinates of M. This is accurate because the zeroed-out regions of the image do not correspond to tissue measurement anymore. As a shorthand, we will denote *T* (*M*_*s*_ ***F***) ∩ *T (M*_*t*_) as *T*_*s+t*_.

We also define a masked average operator *ψ* to average all values of an image *K* over a given set of locations *P*:

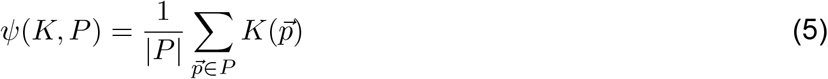

### Elastic Regularizer

For each pixel coordinate 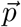, we define a set of neighbors 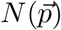 as

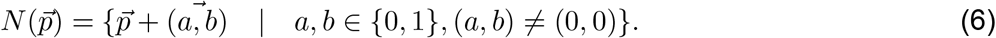

We define elastic energy of a vertex at coordinate 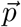 as the sum of elastic energies connected to that vertex. The elastic energy map Ω for a field ***F*** is a mapping between pixel coordinates and the elastic energy of the vertex associated with that coordinate,

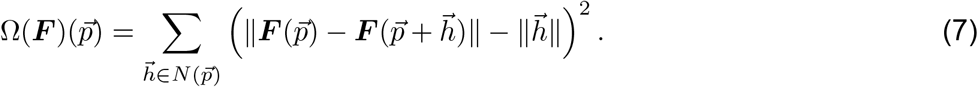

We define elastic regularizer *L*_*er*_ as the masked average of elastic energies for tissue vertices in the source image,

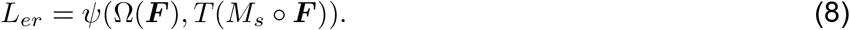

### Image Pair Alignment

For a given resolution, the inputs to an image pair aligner are a source and target image, *M*_*s*_, *M*_*t*_, and the output is a displacement field ***F*** and image encodings *Q*_*s*_ and *Q*_*t*_. The aligner is trained in two stages with two loss functions. During the first training stage, the loss is formulated as combination of similarity between images and elastic regularizer. We define a pixel-wise square error map between images *M*_1_ and *M*_2_ as *E*(*M*_1_, *M*_2_),

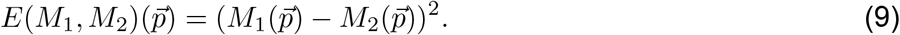

The training loss during the first training stage, *L*_*1*_, is defined as a combination of mean squared error between *M*_*t*_ and *M*_*s*_ *◦* ***F*** masked over *T*_*s+t*_ and the elastic regularizer,

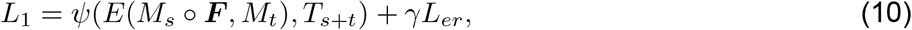

where *γ* is a hyperparameter that balances similarity and deformation loss. In our experiments *γ* varied from 5 to 25, typically with a higher value for lower resolution aligners. The first stage of training proceeds for 100 epochs.

After the first stage of training, the Encoder/Aligner pair is able to produce an initial displacement field, but the image encodings may still contain visual noise. The second stage of training removes visual noise from the image encodings, increases density of the encoding features, and improves the initial displacement field produced by the Aligner.

In the second training stage, the loss is computed as a combination of three components: post-alignment similarity, pre-alignment similarity, and elastic regularizer. On a high level, post-alignment similarity minimizes the difference between encoding after alignment, thus encouraging producing accurate alignment field ***F*** as well as suppressing noise in the encodings.

The post-alignment loss component *L*_*post*_ as mean squared error between *M*_*t*_ and *M*_*s*_ ◦ ***F*** masked over *T*_*s+t*_,

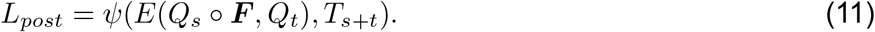

The pre-alignment similarity loss incentives generation of non-trivial encodings by maximizing the encoding difference before the alignment transformation.

In order to increase feature density, we introduce *best sample similarity* error. We first divide the encodings into a set of non-overlapping chunks, *U*. In our experiments, we used square chunks with width and height of 32 pixels. For each chunk, *U*_*i*_, we compute the masked similarity error,

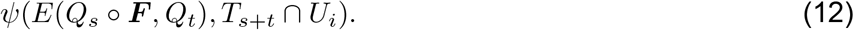

We rank the chunks by their similarity error, and select the bottom k-th percentile of chunks, *V*. We define *best sample similarity* as the average masked similarity error across the set of chunks *V*. The intuition is to select regions of the images that do not contain sufficient feature density to significantly contribute to the pre-alignment similarity loss. In our experiments, we set *k* to be the 50th percentile. Pre-alignment similarity can be formulated as

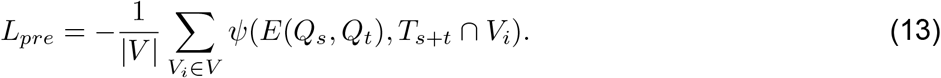

Finally, the stage 2 loss can be formulated as

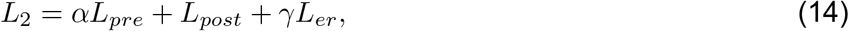

where *α* is a hyperparameter that balances pre-transformation and post-transformation similarity during training. In our experiments α was set to 0.7, and the second training stage lasted for 500 epochs.

After the end of the second training stage, the field is computed for the whole training dataset, and training proceeds to the next scale Aligner/Encoder pair.

### Vector Voting Produces Median Displacements

We introduce vector voting, a function that smoothly computes the median displacement field from a set of fields,

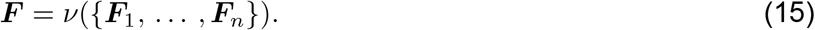

To start, we organize the n input fields into all combinations of subsets that constitute a minimum simple majority, such that

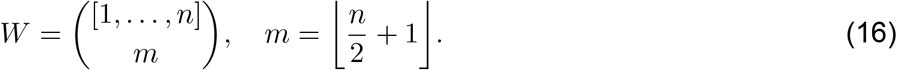

For example, if *n* = 3, then *W* = {{1, 2}, {1, 3}, {2, 3}}. We compute the average similarity between all pairs of fields for each subset, *W*_*k*_ ∈ *W*. In our experiments we used Euclidean distance,

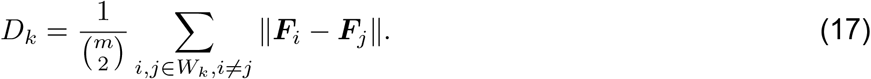

We use a softmin function to map these average similarities to a normalized set of coefficients, such that

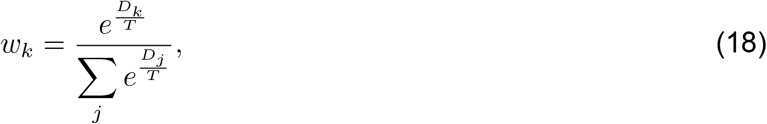

where *T* is the temperature. A lower softmin temperature, *T*, will give outlier vectors less weight in the output, but at a trade-off of potentially including undesirable discontinuities in the field as neighboring positions in the output may be averages from different subsets. For our experiments, we used *T* = 5.7 with *n* = 3. As an added insurance against introducing undesirable discontinuities, the fields may be spatially smoothed when computing their similarity in eq. 17.

For each subset, we distribute its coefficient evenly across its member fields, and the ultimate weight for each field in the input set is the sum of these distributed coefficients. The final field is then

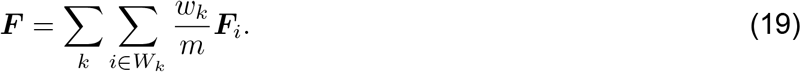

### Vector Voting Corrects Aligner Errors

Given two images *M*_*s*_ and *M*_*t*_ as input, an aligner *ϕ* computes a displacement field

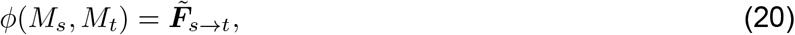

that transforms *M*_*s*_ to be similar to *M*_*t*_. The subscripts *s* and *t* indicate section indices. The tilde sign indicates a field directly produced by an aligner, as opposed to a field that has undergone additional operations.

Aligners can use a previously computed field to transitively align a source to a previous target,

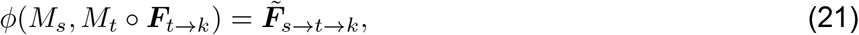

as well as align a transformed source to a target,

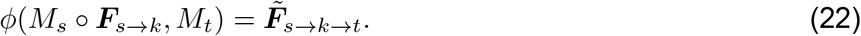

The field, 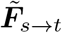, can contain errors if the target image contains missing or corrupted information. If we have a set of images, {*M*_1_, …, *M*_*n*_}, we can make multiple measurements of the same displacement field using transitive alignment. Now we can use vector voting as an error correction procedure with the set of displacement fields,

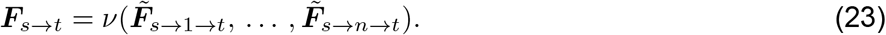

This assumes the Anna Karenina principle: correct transitive fields are relatively similar with other correct fields, while incorrect fields will be different in different ways. The smooth consensus of vector voting preserves desired smooth and non-smooth features of the set of displacement fields.

### Sequential Alignment

With an error correction procedure in place, we can now define sequential alignment. It starts by fixing the first section of the block, so that its field is the identity (section indices relative to the first section),

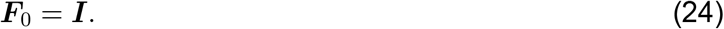

To provide multiple targets for voting on the next few sections in the sequence, we align a few previous sections to the first section,

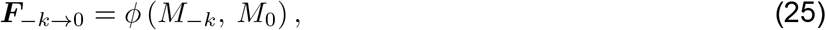

where *k* ∈ {1, …, *n −* 1}. From multiple measurements of the displacement field,

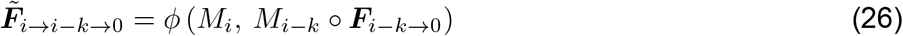

we use voting to produce the final consensus field that aligns a given section to the start of the block,

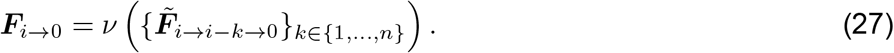

### Block Alignment

Starter sections, *B*, were manually selected by reviewing every 10th section at 1024 nm resolution, and identifying the nearest section that was deemed free of large defects. Block sizes were adjusted accordingly. Section indices are relative to the first section of the entire dataset.

The displacement between a neighboring pair of blocks, or stitching field, was computed by registering the overlapping *n* sections of the two blocks, *S* = { *B*_*i*_ + *k* : *k* ∈ {0, …, *n* − 1}}, then voting over the set of *n* fields that were produced.

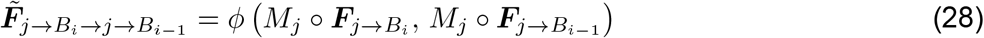

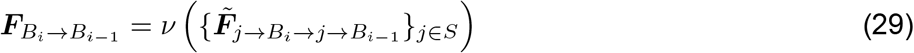

Blocks were stitched together by composing the displacement field of a section computed during block alignment with preceding stitching fields, such that

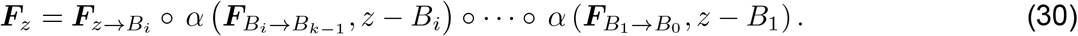

The stitching fields are adjusted, or “decayed”, based on their distance from *z*. For a distance *n*, a stitching field is decayed such that

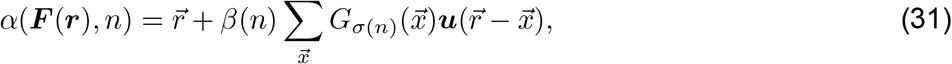

where *G*_*σ*(*n*)_ is a blurring kernel with varying standard deviation that reduces the high-frequency components of the field with distance, and *β*(*n*) is a function that returns a coefficient to dampen the effect of the field with distance. In our experiments, we used

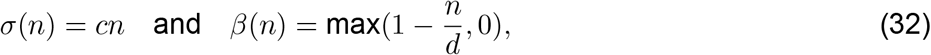

with *c* = 0.2 (measured in pixels) and *d* = 200 (measured in sections). The effect of applying a varying blurring kernel on each stitching field was generated by trilinearly interpolating a MIP hierarchy of the field generated using an iterative 2×2 box filter (Williams, 1983).

### Composition with Decay Preserves Pair Alignment

Block alignment should preserve pair alignment, such that,

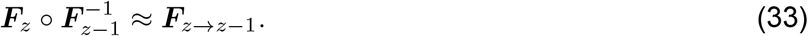

For example, let’s use fields that are in the first block, so that

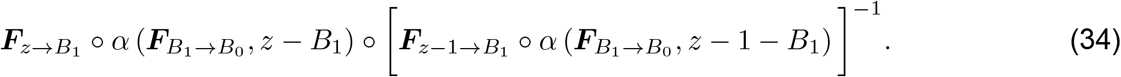

The inverse of a composition is the composition of the inverses reversed, and an inverted aligner field aligns a target to its source,

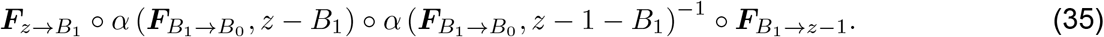

We want to show that the second and third fields are roughly inverses,

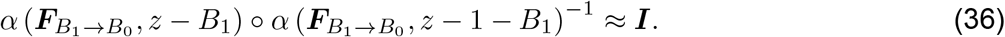

To simplify, let’s set *n* = *z* − *B*_1_. Starting with the vector equation for field decay, eq. 31, let’s define a separate variable for the adjusted displacement,

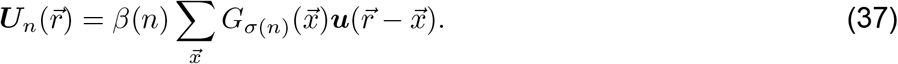

This lets us write the inverse of the adjusted displacements as,

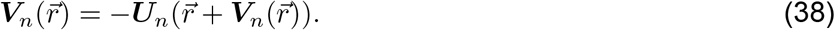

Now we can rewrite eq. 36 as a vector equation,

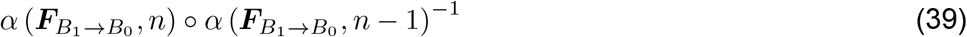

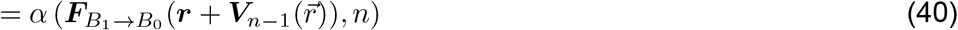

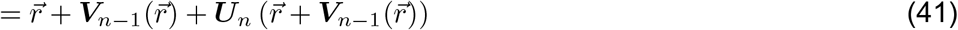

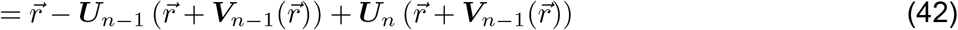

Let 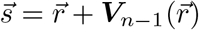, and substitute the definition of the adjusted displacements,

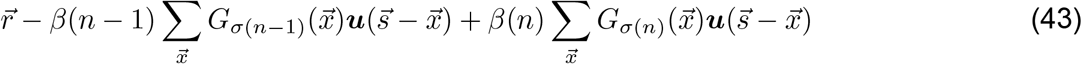

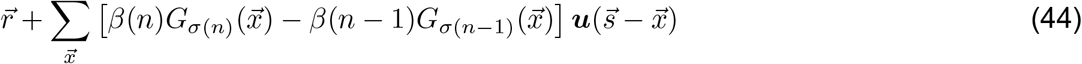

If we use a very large decay distance so that *β(n)* ≈ 1, then the filter convolving the displacements, 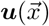, is the difference between two blurring kernels with slightly different standard deviations. The slower that σ increases, the more similar the two blurring kernels, and the closer to producing the identity. Consider using a normalized Gaussian blurring kernel for G with a σ that increases at a rate of 0.02 px per section. This will see that the kernels do not differ to start, increasing to a peak difference of around 5% at about 22 sections, then decreasing to within 1% after 40 sections.

### Large Scale Inference

In order to efficiently apply the proposed approach to large scale datasets, we designed an open source inference framework *corgie* (COnnectomics Registration Generalizable Inference Engine). The main functionalities of *corgie* are handling the dependencies between various steps in the pipeline and handling arbitrarily large images, while maintaining high utilization of distributed workers. Ability to gracefully handle dependencies major is a major difference between *corgie* and other large scale 3D stack processing frameworks(Wu et al., 2021). *corgie* is able to process arbitrarily large images by breaking them into smaller chunks, processing the chunks independently, and assembling the final result from the processed chunk outputs. As quality of convolutional network predictions tends to deteriorate near image boundaries, *corgie* allows the input chunks to overlap, combining their results either through cropping or soft blending. Processing of an individual chunk constitutes an atomic task, which will be assigned to one of the distributed workers. Instead of creating a static computation graph which encodes all of the dependencies between tasks, which may introduce overhead when the number of tasks is large, *corgie* uses dynamic task dispatch. More specifically, each processing step is defined as a Job. Upon being called, a job will yield either a set of tasks or a Barrier dependency, indicating that all of the tasks previously yielded by this job must be completed before the job can proceed.

As an example, alignment of each block will be executed as an independent block-align job. Block-align job is implemented as a combination of other jobs, such as compute-field job, render-job, vector-vote-job, etc. corgie’s task scheduler will distribute tasks from all of the simultaneously running jobs to the same pool of workers, track Barriers and task completion statuses, and schedule new tasks only when the job is ready to proceed. The *corgie* abstraction allows combining these jobs in straightforward, imperative way, while keeping cluster utilization high. *corgie* supports all of the processing steps described in this paper, including image pre-processing (normalization, defect mask computation and burn-in).

### Training Code

The *metroem* package implements the training code necessary for all of the models presented in this work, and is based on the PyTorch deep learning framework. The package is easy to install and operate, allowing users to train new models and finetune existing models on new data without extensive expertise in deep learning. Models trained through *metroem* can be used in *corgie* without any additional conversion, although *corgie* is not limited to using only *metroem* models. Additionally, *metroem* lets the users train aligners for multiple resolution with a single command.

## SUPPLEMENTARY FIGURES

**Fig. S1.**
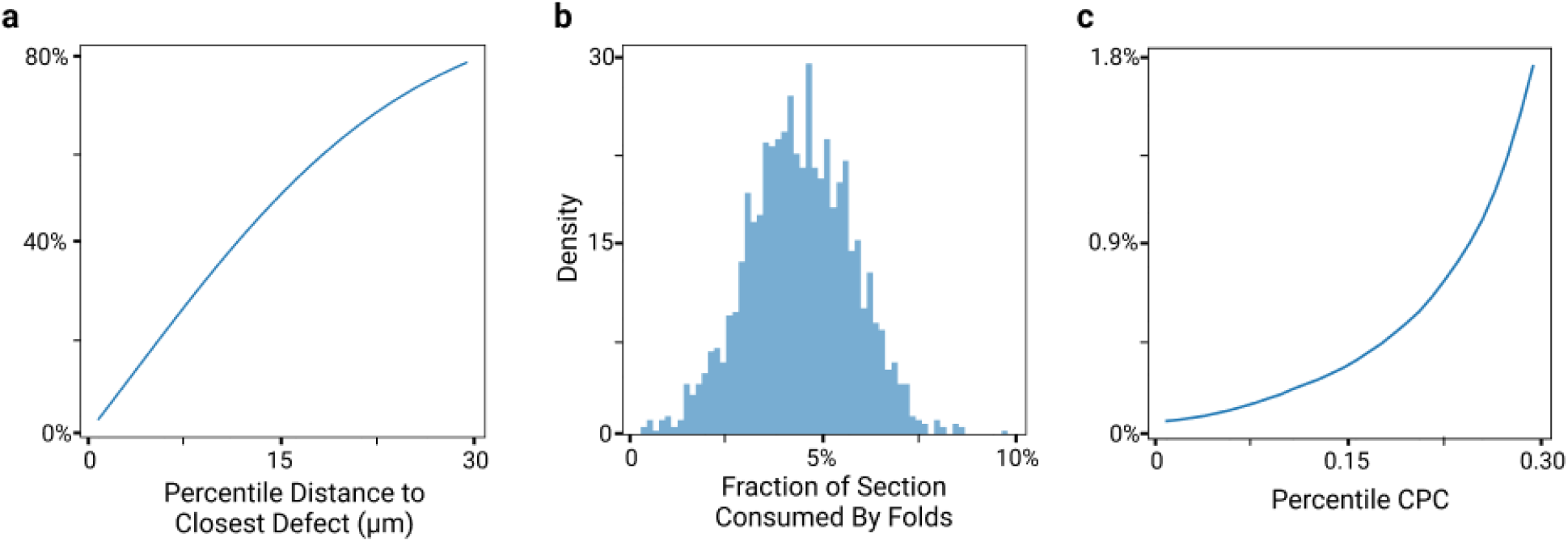
Mouse Cortex Alignment Statistics. Alignment statistics collected over 500 sections of 0.4 × 0.4 mm cutout. The cutout was aligned without vector voting. **a**, Percentile plot for the distance of tissue to the closest defect. Defect locations were identified with a dedicated ConvNet. More than 50% of the tissue lies within 20 μm of the fold. **b**, Per-section distribution of the fraction of tissue consumed by folds. Fold locations were identified with a dedicated ConvNet. On average, 4.6% was consumed. **c**, Percentile CPC plot for each alignment method. CPC is performed on chunks of 2048 × 2048 nm at 64 × 64 nm pixel resolution. Non-tissue chunks, chunks inside blood vessels, cell bodies, and chunks that include a discontinuous artifact were excluded.

**Fig. S2.**
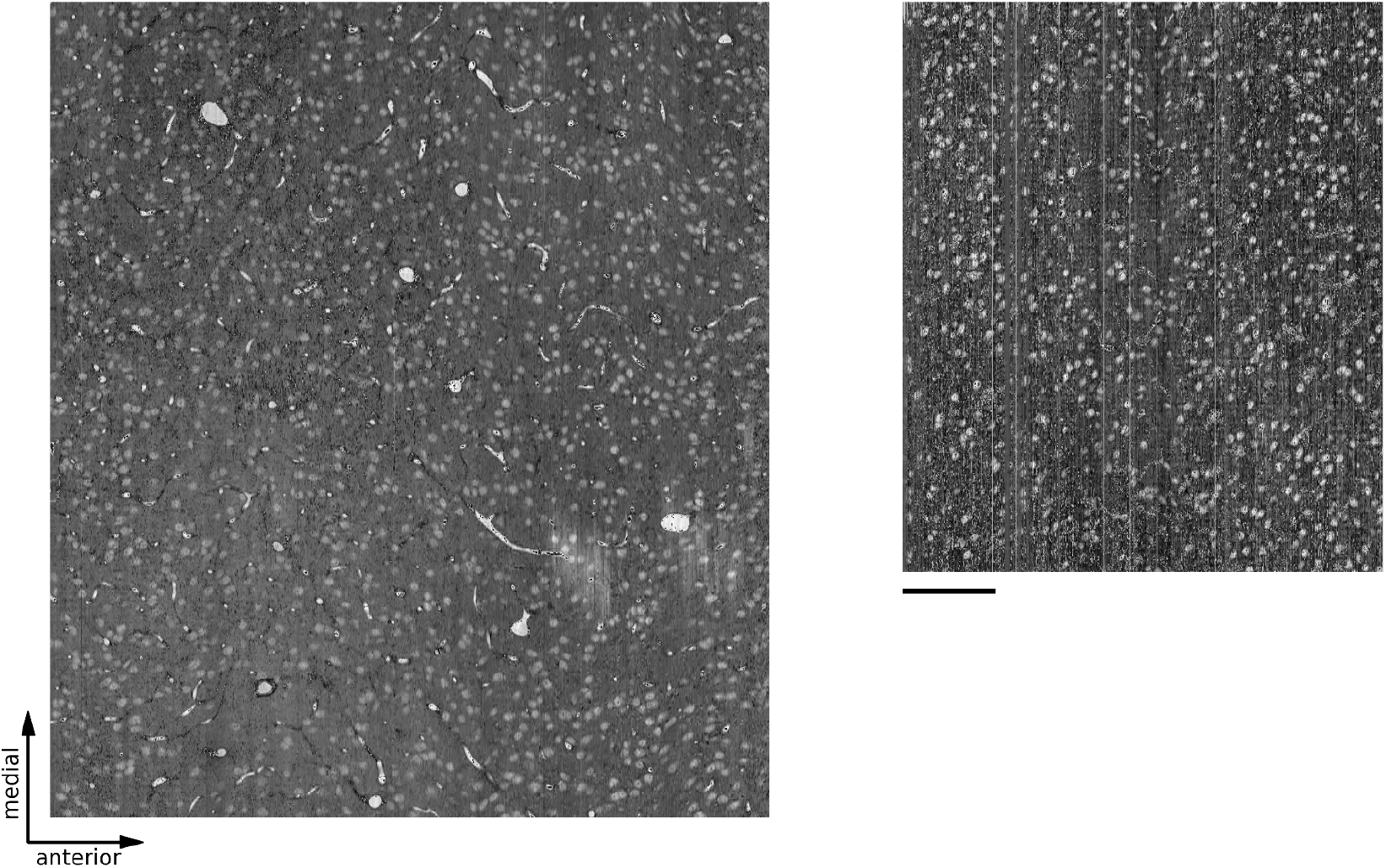
Alignment of mouse cortex datasets with over 10k sections. (left) A view of a cubic millimeter of mouse cortex with 20,000 sections after alignment by our pipeline, where each column represents a section of the data. (right) Similar view of another mouse cortex with 13,000 sections after alignment by our pipeline. Scale bar is 100 μm.

**Fig. S3.**
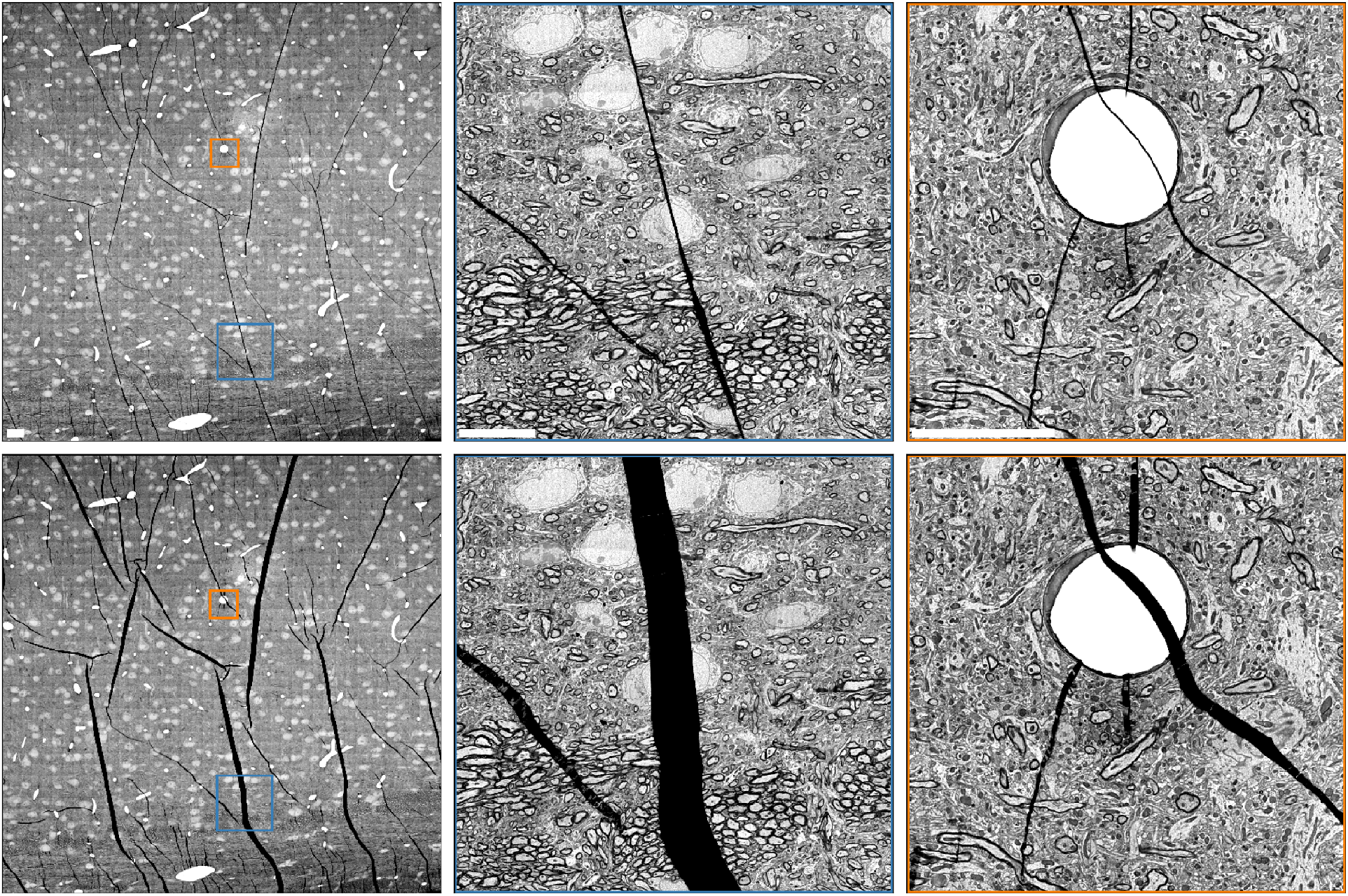
Folds at multiple resolutions. Single section views of mouse cortex data (Fig. S2, left) before (top), and after (bottom) applying our alignment pipeline, with blue and orange insets displayed at higher resolution. Scale bars are 10 µm.

## Footnotes

1 EM Aligner montage diagnostic tool: https://bit.ly/3gI5Cg0

2 Montage SIFT parameter sets: https://bit.ly/3sy44Lf

3 BigFEaTure Aligner (BigFeta): https://github.com/AllenInstitute/BigFeta

4 TrakEM2 solver client: https://bit.ly/3gCuu98

## Notes

https://www.microns-explorer.org/cortical-mm3

https://flywire.ai/

